# The value of prior knowledge in machine learning of complex network systems

**DOI:** 10.1101/094151

**Authors:** David Craft, Dana Ferranti, David Krane

## Abstract

Our overall goal is to develop machine learning approaches based on genomics and other relevant accessible information for use in predicting how a patient will respond to a given proposed drug or treatment. Given the complexity of this problem, we begin by developing, testing, and analyzing learning methods using data from simulated systems, which allows us access to a known ground truth. We examine the benefits of using prior system knowledge and investigate how learning accuracy depends on various system parameters as well as the amount of training data available. The simulations are based on Boolean networks – directed graphs with 0/1 node states and logical node update rules – which are the simplest computational systems that can mimic the dynamic behavior of cellular systems. Boolean networks can be generated and simulated at scale, have complex yet cyclical dynamics, and as such provide a useful framework for developing machine learning algorithms for modular and hierarchical networks such as biological systems in general and cancer in particular. We demonstrate that utilizing prior knowledge (in the form of network connectivity information), without detailed state equations, greatly increases the power of machine learning algorithms to predict network steady state node values (“phenotypes”) and perturbation responses (“drug effects”).

## 1 Introduction and motivation

The ability to better predict the response of a patient, regarding both intended therapeutic effect and potential toxicities, to a candidate drug, radiation, or other treatment modality, would have immediate positive consequences for human health. Currently, cancer patients are usually prescribed drugs based on their tumor type (location, histology, stage). With the advent of targeted therapies, which are designed to interact with biological pathways that are specifically altered in the patient’s cancerous cells, tumor-specific genetic mutations are increasingly being used for drug selection [1]. Examples of gene/disease site pairs for which target therapies exist include HER2/breast cancer, BRAF/melanoma, and EGFR in colorectal and lung. However, even in these target cases, there is a wide spectrum of response to the drugs, which is due to the heterogeneity across patients and within tumors [2, 3].

The problem of predicting how a patient will respond to a particular treatment can be framed as statistical machine learning problem. One can view this as a regression problem if there are quantitative measures of response or a classification problem if the response is binary, e.g. “responders” or “non-responders.” For cancer, which is linked to gene mutations, copy number variations, genomic rearrangements, and epigenetic modifications [4, 5], to perform such a classification, measurements of the genomic state of the patient – from both their healthy tissues and their malignant cells – will serve as key inputs. Supervised machine learning attempts to find statistically valid relationships between inputs and outputs. One of the main barriers to using machine learning for genomics in healthcare is the “large *p* small *n*” problem [6]. The number of genes (*p*) in the human genome is on the order of 20,000, and the amount of additional information above and beyond the gene expression levels of these 20,000 genes is always growing with new assays. Meanwhile, the number of patient samples (*n*) for a cancer trial is typically on the order of hundreds, rarely reaching into the thousands. In situations where *p* >> *n*, patterns will appear in the data by chance (for example, all patients with high expressions of gene *x* do well on the drug) [7]. More generally, it is impossible to find statistically valid relationships without somehow regularizing or compressing the data.

Compressing genomic data into a compact signal could be done without reference to any of the underlying biology of the system that generates the data. However, domain knowledge (biology) could extract more relevant information from the signals and may turn out to be effective, and even crucial, for making high quality clinical predictions. In order to study the question of how prior knowledge can be incorporated into machine learning approaches, we use a computational simulation approach, which allows us access to a known ground truth. More specifically, we generate and simulate networks, analogous to biological systems, and use the data from these simulations to assess machine learning algorithms that predict network behavior with and without the use of knowledge about the underlying system.

## 2 Methods

We use randomly generated Boolean networks [8, 9] as means of producing large datasets to apply machine learning algorithms to. Boolean networks are graphs (nodes and directed arcs) with Boolean logical rules attached to each node. The logical rules update the nodes at each time step. We choose Boolean networks because they have many analogies to biochemical circuits and because they are inexpensive to simulate at large scales (one does not need to solve differential equations, for instance). The entire workflow, from the creation of random Boolean networks, to their simulation, and to the prediction problem, is depicted in Figure 1. Detailed steps are explained in the following sections.

**Figure 1:**
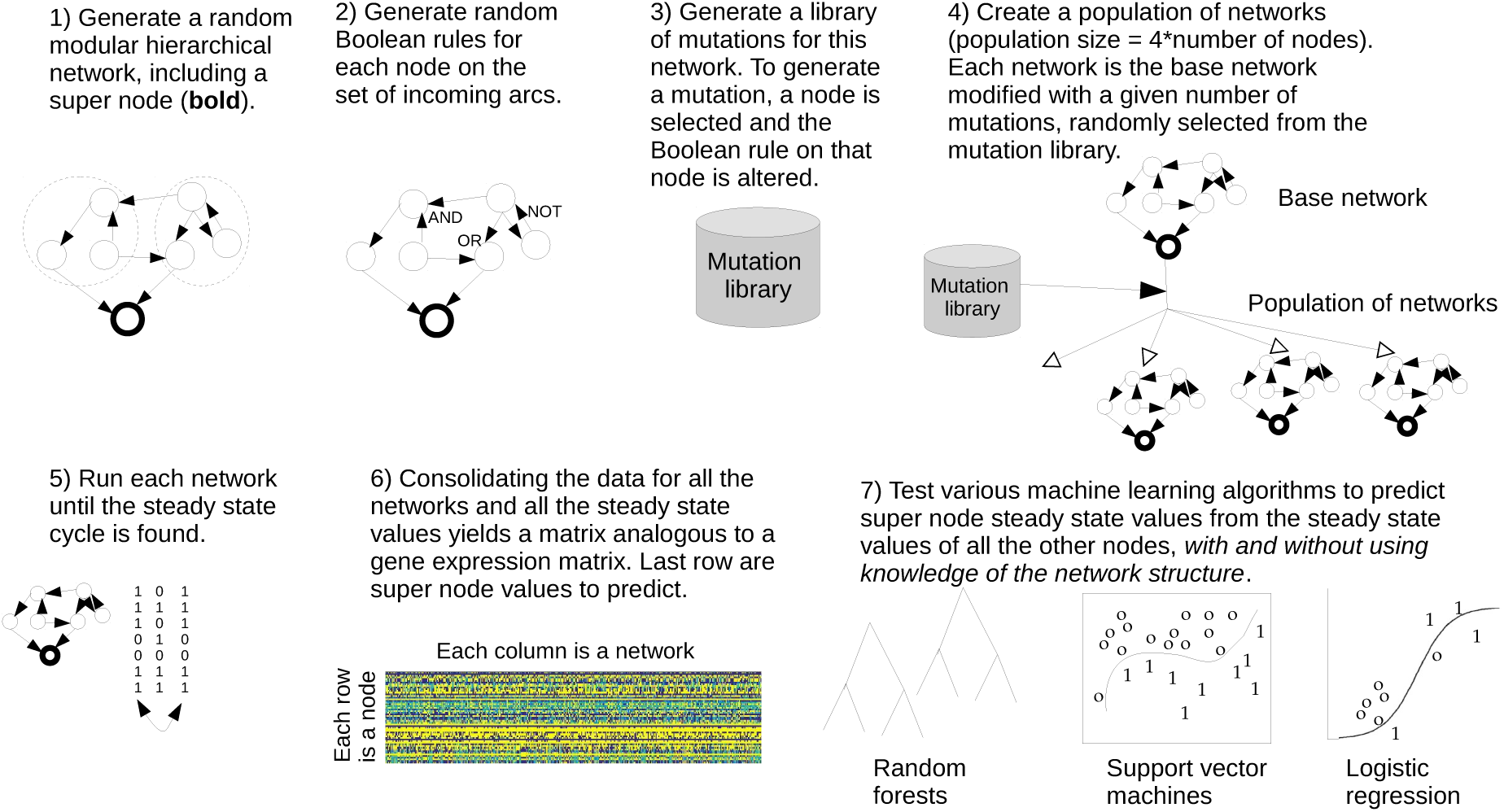
Network generation, simulation, and machine learning problem workflow shown for the phenotype prediction problem (see section 2.3) using random Boolean networks. Further information for Step 1 is detailed in Figure 2.

### 2.1 Random Boolean networks: generation and simulation

The steps for generating a random Boolean network on *N* nodes are 1) Add random directed edges between nodes (we require all networks to be fully connected) and 2) For each node, create a random Boolean rule on the incoming arcs.

For step 1, the graph generation step, we impose a modular and hierarchical structure to the networks in order to mimic the configuration of biological systems [10]. We define a four-leveled hierarchy (roughly: genes, pathways, subfunctions, and functions). We define n_g_ as the number of genes in a pathway, *n_p_* as the number of pathways per subfunction, *n_s_* as the number of subfunctions per function, and *nf* as the number of functions defining the entire system. The total number of nodes (genes) in a network is *N* = *n_f_n_s_n_p_n_g_*. An example of this modular and hierarchical structured network setup is shown in Figure 2. Once the nodes and the module groupings are defined by setting those values, we randomly add edges to the network. For the pathways, we loop through all possible pairs of nodes (directed pairs, meaning pair *a*, *b* is distinct from pair *b*, *a*) and add a directed arc with probability *p_g_* (we do not allow self loops). After making all of the connections within the pathways, we loop through and select all pairs of nodes within the same next level up of the hierarchy, the subfunctions, not including the pairs of nodes that belong to the same pathway (since that pair has already been “visited”). We add a directed arc between two nodes at this level with a smaller probability, *p_p_*. We continue this outward expansion, adding edges between nodes belonging to distinct functions, and then finally across the entire network, with probabilities *p_s_* and *p_f_*, with *p_f_* < *p_s_*< *p_p_*< *p_g_*. Connecting genes in this way mimics the concept of pleiotropy, where a gene (protein) may have multiple functions in an organism.

**Figure 2:**
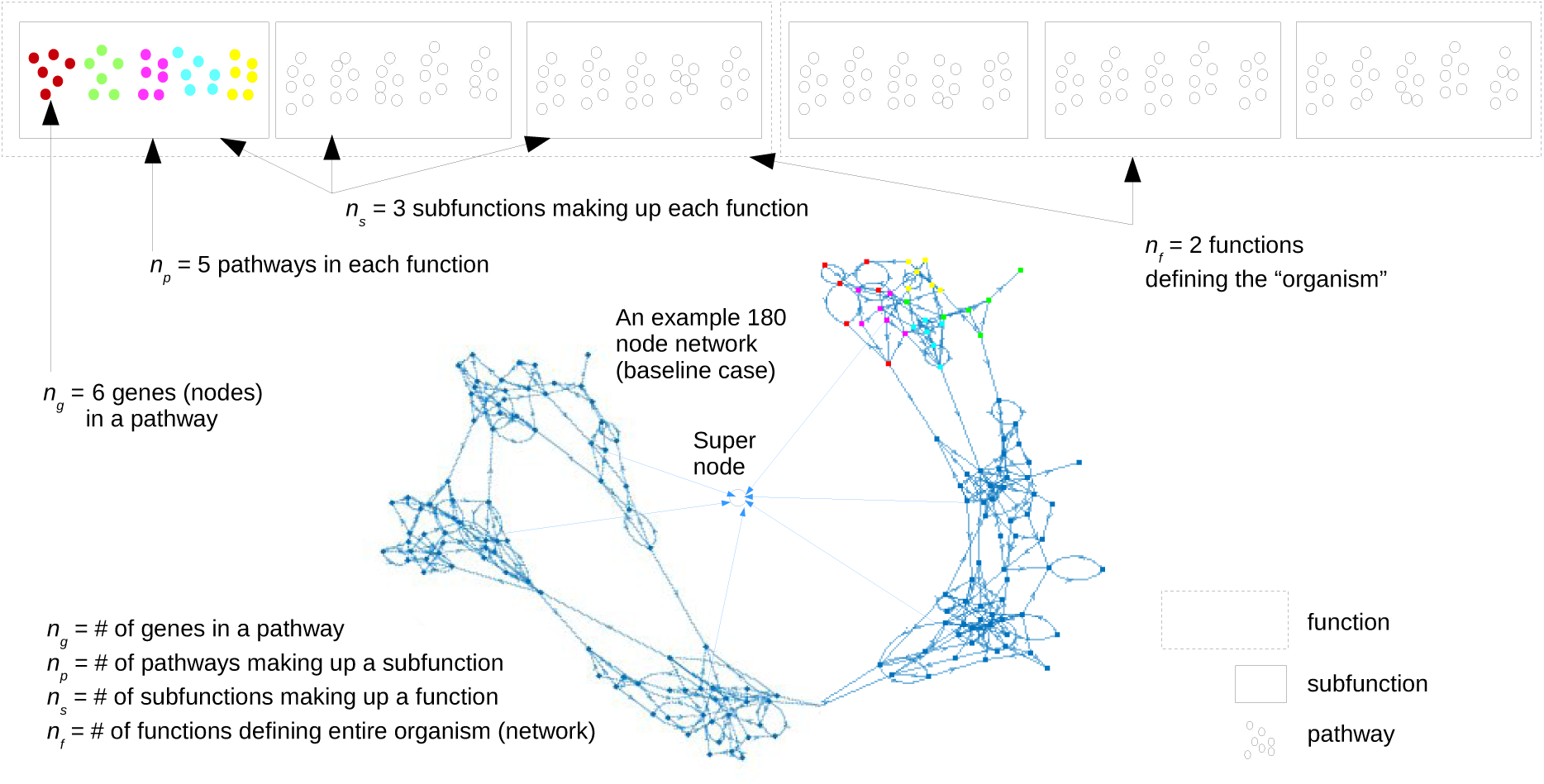
Modular and hierarchical network construction. For the probabilities of an arc between two nodes, we used *p_f_* =.0001, *p_s_* =.002, *p_p_* =.03, and *p_g_* =.28.

We add a special node to the baseline network which we call a super-node, which is meant to be an aggregate assessment of the state of the system. Thus the super-node is a newly created node that has inputs from each different component of the network at a certain chosen level of the hierarchy. For example, if we choose to use the subfunction level of the hierarchy, we select one node from each of the subfunctions to pipe into the super-node, see Figure 2. We choose these pipe-in nodes by selecting a node from each component that only has incoming nodes, with the idea that a node of this type reprents some downstream indication of the state of that component. Since network simulation to steady state (described below) is the most time-consuming part of the data generation step, but including more super-nodes adds on only a negligible amount of simulation time, we add 100 super-nodes to each network. Prior to the learning step, we choose 20 of these 100 supernodes to form 20 distinct learning problems from each full population simulation (see 2.4 for details on how the 20 supernodes are chosen).

Next we add random Boolean logic rules to each node, including the super-nodes. Nodes may have an arbitrary number of incoming arcs. If a node has zero incoming arcs, then that node’s initial condition is the value that persists at this node. If a node has a single incoming arc, then there are only two possibilites for a Boolean function: identity or negation. Thus for these nodes we flip a coin to choose which type we put on that node. For nodes with two inputs, we have many more options. Either input can be negated, and for combining the (possibly negated) inputs we can choose AND, OR, or XOR, and we allow the option to negate the final result as well. For nodes with *K* > 2 incoming arcs, we randomly pair up the arcs and each pair gets combined with a random Boolean function (there will be one triplet if *K* is odd, for which we construct a 3-way Boolean). We then take these 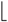*K/*2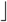 functions (*f*_1_, *f*_2_,…, *f*_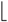_*_K/_*_2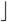_) and combine them into a single Boolean function as follows. We join the last pair of functions (*f*_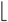_*_K_*_/2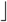–1_, *f*_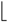_*_K_*_/2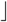_) into a Boolean function, creating *F*_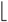K_*_/_*_2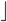_. We then expand this function by adding on the remaining *f* functions one at a time: *F*_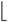K_*_/_*_2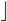_ is combined with *f*_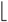_*_K_*_/2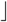-2_, this group is combined with *f*_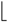_*_K_*_/2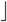-3_, etc. until all of the functions have been used. An example for a node with four incoming arcs is shown in Figure 3.

**Figure 3:**
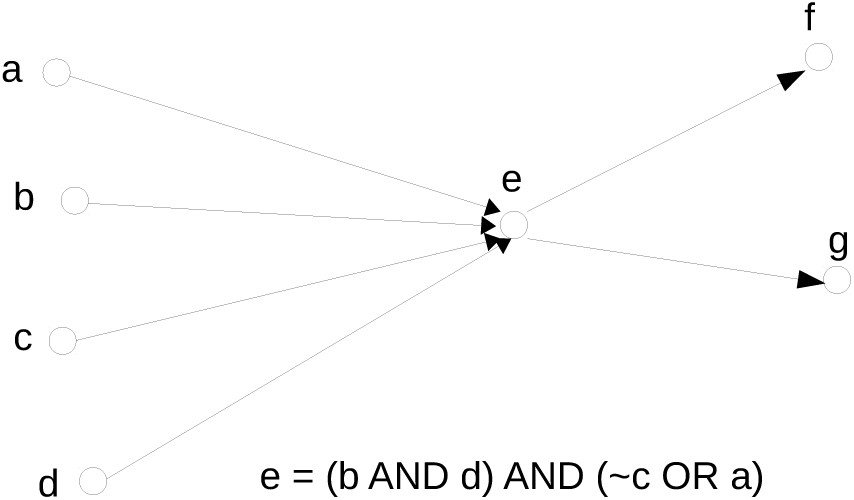
Iterative pairwise construction of a compound Boolean logical statement. In this example, the first two input nodes randomly chosen were **b** and **d**, which got connected with an **AND** statement. Next, node **a** was combined using an OR statement with node **c**, which got negated. Finally, these two pairs are joined together with an AND statement.

With the Boolean logic in place, a network can be simulated. To begin a simulation, nodes are randomly intialized to either 0 or 1. At each time step and at each node, the node’s next value is computed based on the current incoming node values and the Boolean statement. For each node, the result of the Boolean logic gets “broadcast” to all of the nodes connected to that node via its outgoing arcs. Thus, nodal values are associated with the nodes and the outgoing arcs of each node. We use synchronous updating, meaning each node is updated simultaneously at each time step. For the update, each node uses the incoming state values and its logical rule to update its own state value.

Since the networks have a finite number of nodes *N* and each node is either in state 0 or 1, there are a finite (2*^N^*) number of possible network states and therefore at some time the state will repeat. When such a cycle is detected, the simulation is terminated and the node state values across the cycle (the sequence of 0s and 1s that the node goes through) are averaged to produce a steady state nodal vector. If a cycle is not found after some large number of state transitions, we terminate the search and replace that member of the population with a new member (a new set of mutations).

### 2.2 Random Boolean networks: population creation

To create a population of members (which could represent cell lines, patient tumor samples, etc.) we start by creating a baseline Boolean network as described above and then, for each member of the population, we take this baseline network and put random mutations on it, thus producing a diversity of related networks. Mutations come in the following forms:

1. If a node has no predecessors, it is mutated by changing (i.e. negating) its initial condition.
2. If a node has only a single predecessor, it is mutated by negating the rule on the incoming arc.
3. If a node has two or more predecessors then the following mutations are possible:

- The node is activated, meaning independent of the inputs the output is always 1.
- The node is deactivated (output always 0).
- Logic change: one of the symbols in the Boolean algebra is changed. AND gets changed to OR or XOR, etc., or an entity is negated.

To create a population, we first create a library of *L* mutations. For each *i* ∈ 1,2,…, *L* we select a node from the network, with replacement, and create a mutation from the above mutation type list. This serves as our mutation library. We do not allow the pipe-in nodes nor the supernodes to be mutated. For each member of the population, we randomly select a smaller number of mutations *R* from the mutation library and apply those mutations (i.e. we overwrite the Boolean logic or initial conditions) to the baseline network. Once a population is created, each member is simulated until a steady state cycle is found. The steady state values are used as data for the machine learning problem.

### 2.3 Two learning problems: phenotype prediction and drug effect (perturbation) prediction

We define two learning scenarios for our network datasets. In the *phenotype prediction problem* we attempt to predict the steady state value of the super-node based on the steady state value of the other nodes in the network. In the *drug prediction problem* we simulate a more clinically relevant scenario. We first simulate the patients in a population and record their steady state values on all nodes except the super-nodes. This data represents the pre-treatment patient assay, for example a gene expression profiling of their tumor. We then simulate a drug added to each of the patients by applying the same randomly chosen mutation to each of patient networks (which mimics the basic understanding of a targeted agent, where a certain gene is affected in a consistent way across the patients). We then simulate all of the patients again with this mutation and record the steady state value of the super-node. For patients in the test set, we then try to predict the post-drug super-node value from the pre-drug network steady state values. This *drug prediction problem* is thus once removed and leads to a harder learning problem.

For all problems in this work, we use classification algorithms, which requires rounding steady state values of the super-nodes to 0 or 1. We verified that random forest and support vector machines, the best performing algorithms, are also effective in regression mode for this problem, but we focus on classification, for simplicity, and do not show any regression results.

For both the phenotype and the drug effect prediction problem we also investigate an idealized learning problem where the node value to predict is a Boolean function of the steady state values of the other nodes (in our normal case, super-node values are based on the full network dynamics). This yields an easier “pure Boolean logic” learning problem, where we take the steady state node values, round them to 0 or 1, then apply the randomly generated Boolean logic statement to those values to form the super-node value. This problem assess the capability of the learning algorithms to learn pure Boolean functions.

### 2.4 Description of baseline network and datasets

The parameters for the baseline network are as follows: *n* = [*n*_*g*_, *n*_*p*_, *n*_*s*_, *n_f_*] = [6, 5, 3, 2] and *p* = [*p*_*g*_, *p*_*p*_, *p*_*s*_, *p*_*f*_] = [.28,.03,.002,.0001], see Figure 2. Pipe-in nodes are selected at the subfunction level, thus there are typically 6 (3 × 2) pipe-in nodes out of the total of *N* = 180 nodes. There will be fewer than 6 pipe-in nodes if one or more of the subfunctions had no nodes with only incoming arcs. For the baseline case, the mutation library size is *L* = 180 and the number of mutations imposed on each network sample is *R* = 36 (20% of the number of nodes).

For each set of network parameters (*n, p, L, R*) studied, including the baseline set, we generate five network topologies (directed arc connections) and for each of these network topologies we generate five different Boolean rule sets, for a total of 25 distinct networks per parameter set studied. For each of these 25 base networks, we generate 4N samples by drawing random mutations from the mutation library and applying them to the base network. Thus for the baseline case, where *N* = 180, we generate and simulate to steady state 720 networks. For each network we create 100 super nodes to test prediction algorithms on. After the 4*N* simulations we choose the 20 supernodes that have the greatest variation among the samples (that is, we choose the supernodes that are closest to having a 50-50 split between samples of phenotype 0 or 1).

### 2.5 Machine learning with and without prior knowledge for various network parameters

We test the following machine learning algorithms: logistic regression, lasso and elastic net regularized logistic regression, support vector machine (SVM), random forest (RF), principle component analysis (PCA) compression with RF, and nearest cluster. The regularization parameter λ for lasso and elastic net is estimated as the largest value in a given sequence that gives a non-null model (a null model has all the *β* terms equal to 0) and the deviance (the model fit) is estimated using 10-fold cross validation on the training set. For SVM, a quadratic kernel consistently outperformed the linear and radial basis function kernels and this is subsequently used in our results. For RF, we use an ensemble of 75 decision trees and each tree used a random 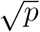 predictors to make splits on, where *p* is the number of nodes seen by the algorithm.

For no prior knowledge, we apply the algorithms to the complete steady state data from all the nodes in the networks (except for the super-nodes, which are the ones we try to predict). To demonstrate the use of prior knowledge we apply machine learning algorithms using the nodes that are suspected, based on network connectivity, to be better predictors of the super-node values. The simplest possibility, and the one we use, is to use the nodes that are directly connected to the super-nodes, the pipe-in nodes. Additionally, we examine the use of patient-specific mutation information, either alone or along with the steady state node values, in the prediction algorithms.

We generate and run many sets of experiments to understand how classification accuracy depends on various network parameters, including network size, number of pipe-in nodes, mutation rate and mutation library size, all with and without the use of prior knowledge.

### 2.6 Univariate selection of nodes to include in the machine learning

In order to improve on “no prior knowledge” learning method, but still without utilizing prior knowledge, we study a few versions of pre-selecting nodes from the entire set based on how well they separate the data in a univariate mode. Samples are divided into two classes based on their thresholded supernode value (either a 0 or 1). Then for these two groups, a t-test was conducted for each node to assess if the node had any predictive power, i.e. if the steady state node value was significantly different for the two groups. P-values for each node were ranked by importance (lowest p-value to highest p-value). In order to find the optimal set of nodes, we create a classification model (e.g., random forest) using the most important node, then the two most important nodes, etc. until we use all of the nodes. These models are then used on the test data and the optimal classifier from all of these *N* models is used as our result, which represents the best we can do for this style of univariate node selection.

We also run the univariate node selection strategy replacing the t-test with a mutual information score as well as a chi-squared score, which performed the same as the t-test so we suppress these results.

### 2.7 Data and code availability

We make available a large set of data from this paper as well as the code base (in Matlab, version R2015b) to generate and analyze new datasets. We used native Matlab functions for all machine learning algorithms. For code and data see https://gray.mgh.harvard.edu/people-directory/71-david-craft-phd.

## 3 Results

We begin by assessing a variety of commonly used machine learning algorithms on our baseline phenotype prediction case. Random forest, support vector machine, logistic regression, lasso (which was similar to elastic net, so results not shown), and nearest cluster, are compared in Figure 4a, which clearly shows the advantage of RF and SVM. We select RF as our baseline learning algorithms going forward. Figure 4a also displays the key result that classification accuracy improves when the algorithms use information from only the pipe-in nodes instead of all of the nodes. This is a direct demonstration of the value of prior knowledge, a consistent theme throughout our results. The p-value for this comparison, from a paired t-test, is <.001. We get similarly small p-values for every comparison of prior knowledge vs. no prior knowledge, and also for the comparisons between any two algorithms, thus we do not continually report p-values. In fact, for all comparisons between two groups, whether paired (when the results come from the same set of networks) or unpaired, we achieve significant differences due to our large sample sizes, 500, even when the differences are not practically significant. For this reason, we opt to report *effect size*, which incorporates the magnitude of the difference in means (mean classification accuracy in our case) of the populations [11]. We opt for the specific version of effect size called Common Language Effect Size (CLES) [12]. CLES gives the probability that a random draw from one group will exceed a random draw from the other group, and is defined for non-paired group testing. For paired tests, we do not report the vanishingly small p-values throughout; for unpaired testing, we report the CLES probabilities.

**Figure 4:**
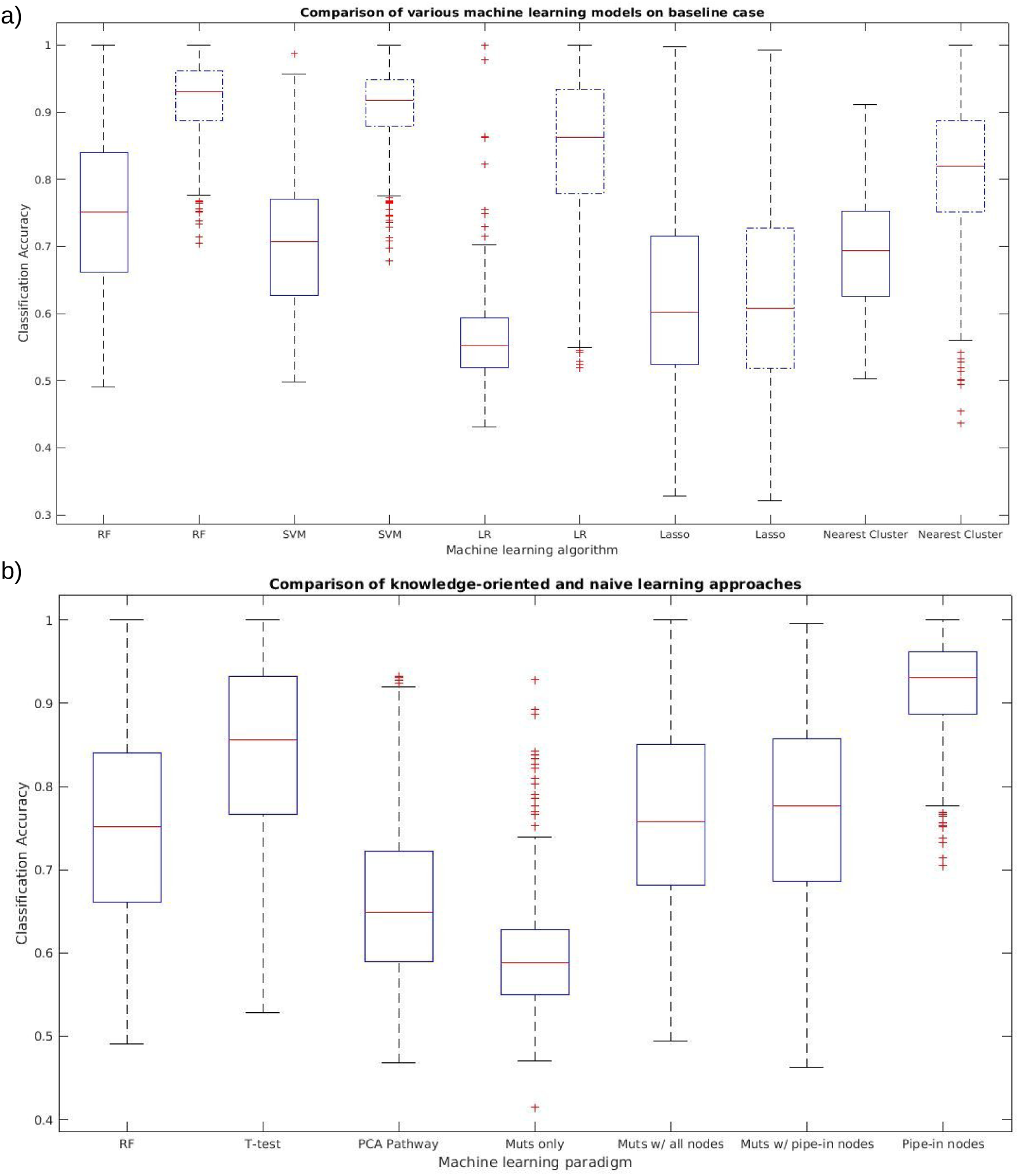
Algorithm comparison on the set of 25 baseline 180-node networks, each with 20 distinct supernodes (20 prediction problems per network) for a total of 500 learning problems. Each dataset contains 720 samples. (a) Box-whisker plots for classification accuracy for various algorithms for all nodes (solid) and just pipe-in nodes (dashed). (b) Using RF, we compare no prior knowledge to various types of prior knowledge: no prior knowledge baseline, no prior knowledge but using t-test on the training set data to identify predictive nodes and then using just those nodes in the random forest algorithm, prior knowledge in the form of PCA compression of the data from each subfunction module (which requires network connectivity knowledge) into two principle components, mutation knowledge only, mutation knowledge in addition to baseline steady-state data from the 180 nodes, mutation knowledge plus steady-state data from just pipe-in nodes, and the pipe-in node result. For visual comparison, the first and last box-whisker plots are repeats from the first two box-whisker plots in (a).

Figure 4b displays that node selection by t-test (similar results for chi-square and mutual information univariate node selection, results not shown) improves the “no-prior knowledge” learning technique, but not as much as prior knowledge node selection (more results of the t-test selection method are discussed below). PCA compression of the subfunction node sets steady state values apparently results in a loss of useful information and performs poorly. We also see that using binary information of which mutations a patient has, while offering some predictive value by itself, adds noise and thus weakens the performance of the learning algorithms when node steady state information is available. Similarly, using additional network connectivity knowledge of which nodes are directly connected to the pipe-in nodes weakens the strong signal of the pipe-in nodes and thus makes classification accuracy drop (results not shown).

We explore the t-test node selection strategy in more detail in Figure 5. In 5(a), the solid curve shows that there is an optimal number of nodes to use when using a t-test selection strategy, in this case around 18. The dashed curve shows that the t-test only gradually picks out the pipe-in nodes. If the t-test idea worked perfectly, the pipe-in node would always be the top ranked by the t-test, but we see that there is a long tail where, if the pipe-in node is not ranked in the top few nodes, than it could appear anywhere in the ranked order, Figure 5b.

**Figure 5:**
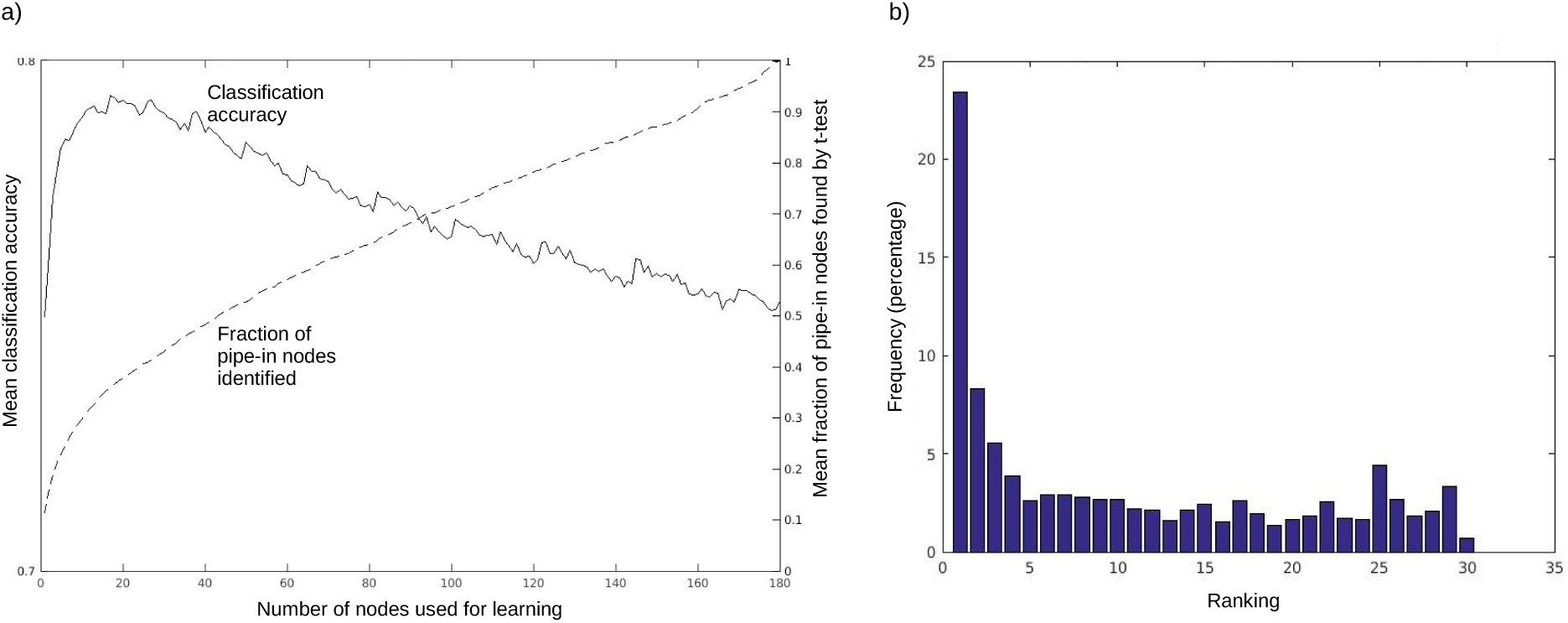
Selecting nodes to use based on a univariate t-test. Nodes are sorted based on their p-value from the t-test which assesses how useful they are in the classification problem. The solid line in a) shows the overall accuracy of the random forest algorithm as nodes are successively added in from the sorted order, accuracy peaking around 18 nodes. The dashed line shows the average number of pipe-in nodes that have been selected by the t-test method for the given number of total nodes added. The linear increase of this curve until it reaches 1 demonstrates that the t-test method does not reliably uncover all of the pipe-in nodes, which is further emphasized in b), a histogram which shows at the node set sub-function level (the place in the hierarchy for our baseline network where pipe-in nodes are selected from) what ranking the pipe-in nodes are in the t-test: less than 25% of the time a pipe-in node is the top ranking node from the relevant node set.

Figure 6 demonstrates the correlation between area under the receiver operating characteristic curve (AUC), a standard measure of performance of classification algorithms, and classification accuracy (CA, which is fraction of super-nodes classified correctly). Since they correlate well and since CA is cheaper to compute, we use CA throughout to compare algorithms.

With RF and CA established as our learning algorithm and performance measure we look at the phenotype and the drug effect prediction problems, for full dynamic network generated data and for datasets generated from the steady state nodal values. These results, shown in Table 1, verify that the drug effect prediction problem is slightly harder, and if the underlying Boolean function is based on steady state data rather than the network dynamics data, the problem is slightly easier. Noteably, all problems are of overall similar difficulty. We choose the phenotype prediction problem with actual network dynamics as our baseline problem type to investigate further.

**Figure 6:**
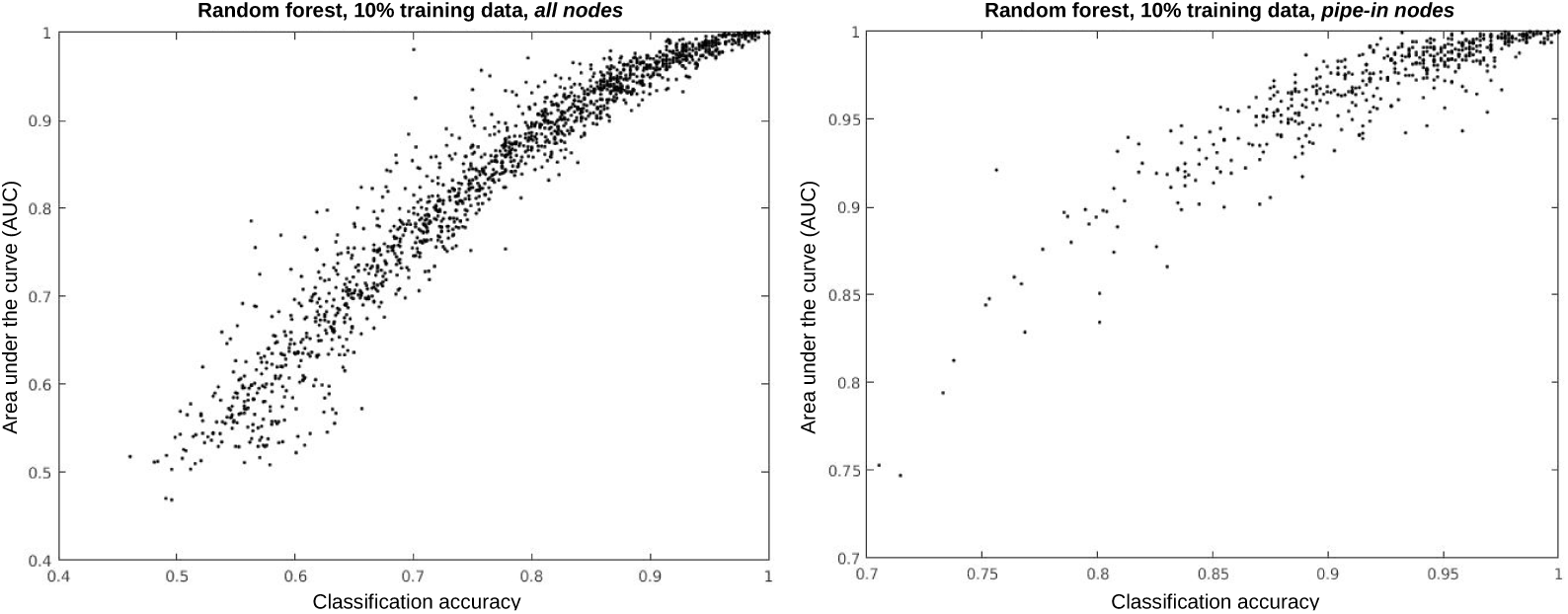
Correlation of AUC and classification accuracy for 500 learning problems of the baseline 180 node networks for (a) all nodes and (b) pipe-in nodes (prior knowledge).

**Table 1:** Four learning problems compared using classification accuracy [mean (standard deviation)]. Phenotype prediction attempts to predict the steady state value of a single network node based on access to the steady state values of the other nodes. The drug effect problem is the two step network simulation and prediction problem as described in the text. The right column repeats these experiments for an idealized setting where the node values to predict are functions of the steady state results rather than functions of the actual system dynamics. The steady state idealization is meant to test the power of the the random forest algorithm to uncover pure but complex Boolean logical rules.

|  |  | Full dynamics | Steady state |
| --- | --- | --- | --- |
| Phenotype | All | 0.80 (.11) | 0.81 (0.12) |
|  | pipe-ins | 0.94 (0.05) | 0.97 (0.03) |
| Drug effect | All | 0.79 (0.11) | 0.80 (0.12) |
|  | Pipe-Ins | 0.92 (0.07) | 0.94 (0.06) |

Table 2 summarizes how the learning problem depends on various network parameters. For all results, we compare no prior knowledge (“all nodes”) to prior knowledge (“pipe-in nodes”) and see that using prior network knowledge consistently wins over no knowledge. Network size in our setting does not greatly affect the difficulty of the learning problem. CLES for network sizes of 90 nodes vs 180 nodes is 64%. Thus, 64% of the time a randomly selected 90 node network dataset has a higher CA than a 180 network dataset (this value is the same for all nodes and for pipe-in nodes). The 180 node networks compared to the 800 node networks are even less distinguishable, with CLES = 60%.

As the number of pipe-in nodes increases from 2 to 6, the learning problem gets more difficult (CLES = 70% for all nodes, 85% for pipe-in nodes learning), as expected. From 6 to 30 pipe-in nodes has much less an effect, with CLES 55% and 59% for all and pipe-in, respectively.

**Table 2:** Classification accuracy [mean (standard deviation)] of random forest algorithm for various network populations. For ease of comparison, the baseline case (·) is repeated for each bundle. For network size runs, the number in brackets is the number of pipe-in nodes.

| Experiment |  | All nodes | Pipe-in nodes |
| --- | --- | --- | --- |
| Network size | 90 [6] | 0.81 (0.13) | 0.94 (0.06) |
|  | · 180 [6] | 0.75 (0.11) | 0.92 (0.05) |
|  | 800 [16] | 0.79 (0.11) | 0.94 (0.05) |
| Number of pipe-in nodes | 2 | 0.84 (0.14) | 0.98 (0.02) |
|  | · 6 | 0.75 (0.11) | 0.92 (0.05) |
|  | 30 | 0.77 (0.12) | 0.89 (0.08) |
| Mutation rate | 10% | 0.84 (0.11) | 0.94 (0.06) |
|  | · 20% | 0.75 (0.11) | 0.92 (0.05) |
|  | 40% | 0.81 (0.10) | 0.95 (0.05) |
| Size of mutation library | 90 | 0.86 (0.11) | 0.96 (0.04) |
|  | · 180 | 0.75 (0.11) | 0.92 (0.05) |
|  | 360 | 0.78 (0.12) | 0.93 (0.06) |

In the limit as the mutation rate approaches 0, all the members of the population would be the same thus leading to a trivial learning problem. However, regarding mutation rate and mutation library size, we do not see large effects. The only CLES scores for mutation rate that exceed 70% are for all nodes, mutation rate of 10% vs 20% (20% mutation rate is harder to classify). For mutation library size *L*, only the jump from *L* = 90 to *L* = 180 gives a CLES over 70%: both all and pipe-in nodes learning give CLES = 76%. Thus in all cases where we expect the learning problem to get more difficult with increasing (network size, number of pipe-in nodes, mutation rate, and mutation library size) we see a stronger effect initially but a saturation of difficulty to somewhere between 75% and 80%.

Figure 7 shows how classification accuracy increases with training set size (this is all paired data, all results p-values <.001), for both uninformed learning and prior knowledge learning. In particular, for this dataset we see that the uninformed learning requires a dataset size of 50% of our full patient sample (360 samples out of 720) to reach the same accuracy as prior knowledge-based learning reaches with only 5% (36 samples) training data size.

In order to compare further the value of additional training data versus the value of prior knowledge, we run RF on the 500 baseline datasets for training set sizes from *i* = 10% (72 samples) up to 90% in increments of 10% assuming no prior knowledge (i.e. using all the network nodes). To mimic partial prior knowledge, we assume we know only *j* of the pipe-in nodes, and vary *j* from 1 to all of them, which is 6 for the baseline networks. For each *j* ≤ 6 and for each learning run, we randomly select *j* of the 6 actual pipe-in nodes to perform the learning problem with. For this prior knowledge investigation, we assume we are training on 10% of the data (the default training set size). For each *i, j* pair, we can compute the average classification accuracy for the no prior knowledge *i*th run and the prior knowledge *j*th run, and form the difference CA(*j*) - CA(*i*). Plotting these differences as a 2D matrix gives us a visual comparison of the value of each type of information, see Figure 8. The top left of this figure demonstrates that prior knowledge with a 10% training set size dominates the no prior knowledge machine learning of up to 50% training size. We see a sharp drop in classification accuracy as prior knowledge, in this case the number of pipe-in nodes that are known, decreases. This explains why the t-test method, Figure 5, while improving on the naïve no prior knowledge method, is still inferior to full knowledge of the pipe-in nodes: the t-test does not reliably select out the pipe-in nodes, and not having all the pipe-in nodes when learning with only 10% of the data, as seen in Figure 8, is greatly detrimental.

## 4 Discussion and conclusions

Since genomic data has become widely – and increasingly cheaply – available there has been much effort to analyze it to discover (or, as is often the case, rediscover) biological mechanisms and to advance medical practice. More often than not, genomic data is analyzed as an independent data problem, and no inputs from known biology are used to regularize the data. A common result of such papers is “our methods uncover the genes *x* and *y* that are known to be important in this context, but also reveal gene *z*, which has not previously been implicated in this setting.” This type of analysis begs the question: if we already knew that genes *x* and *y* were important, and they were somehow put into the method up front, perhaps more information could be extracted from the data. Stated another way, if we use known biology, rather than ignoring it, perhaps we can increase the power of our machine learning methods. Statistical machine learning should complement findings from wet-lab biology, not compete with it.

We demonstrate for idealized systems loosely analogous to biological systems that incorporating prior knowledge greatly increases the ability of machine learning algorithms to make predictions of the system behavior. This indicates that it will be prudent to investigate how to incorporate biological knowledge into machine learning algorithms. Our work suggests that identifying signals (e.g. proteins) that are downstream in a pathway, and offer an indication of whether or not the pathway is functioning, could be a more useful signal for the machine learning than the complete set of signals. However, we stress that our work is suggestive rather than immediately practically applicable, and which are the best signals to focus on for particular biological contexts remains to be investigated.

**Figure 7:**
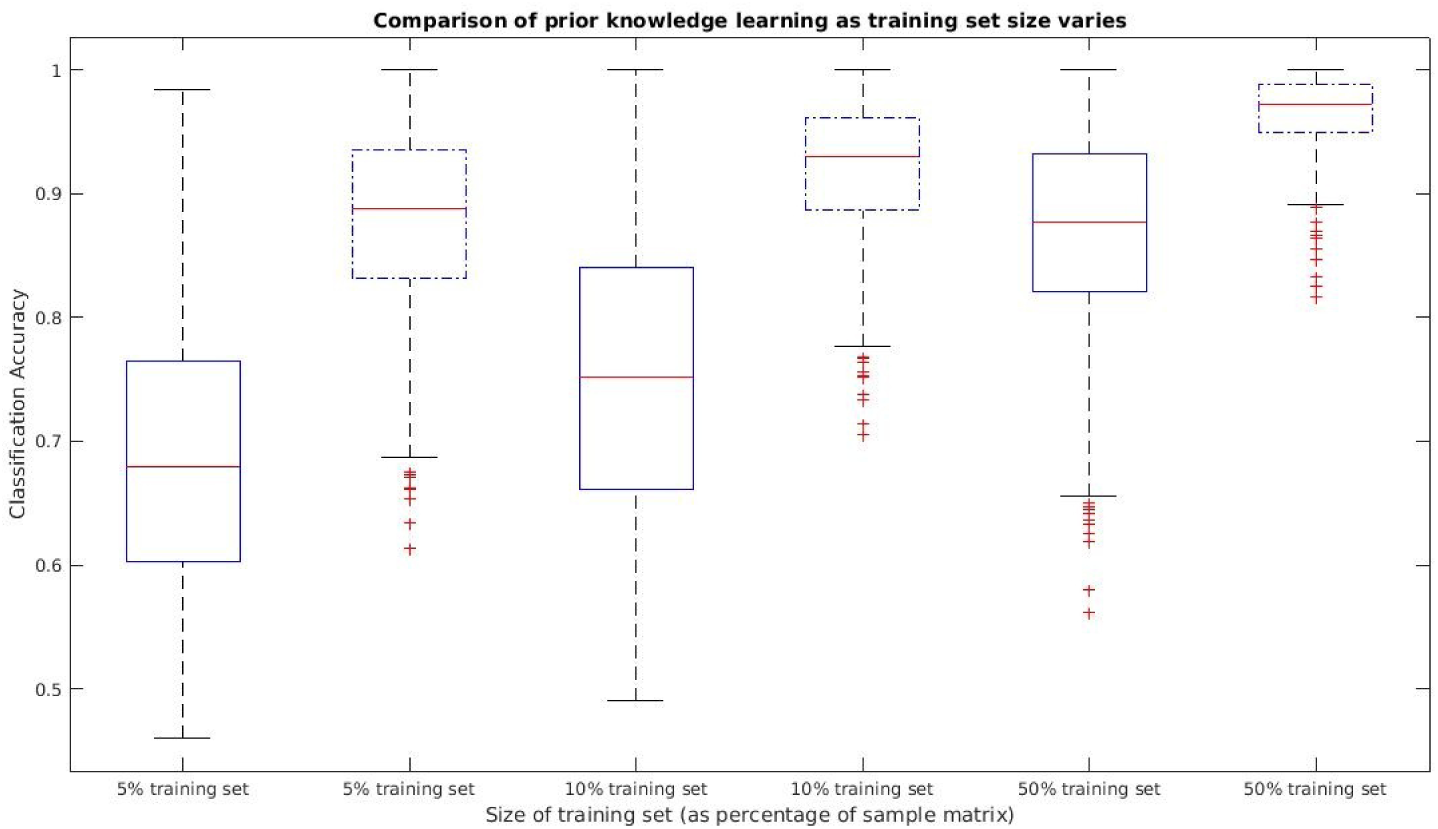
Classification accuracy for the baseline networks for various sizes of the training set. Sizes are percentages of the number of samples we generate in total (720 for the baseline networks).

**Figure 8:**
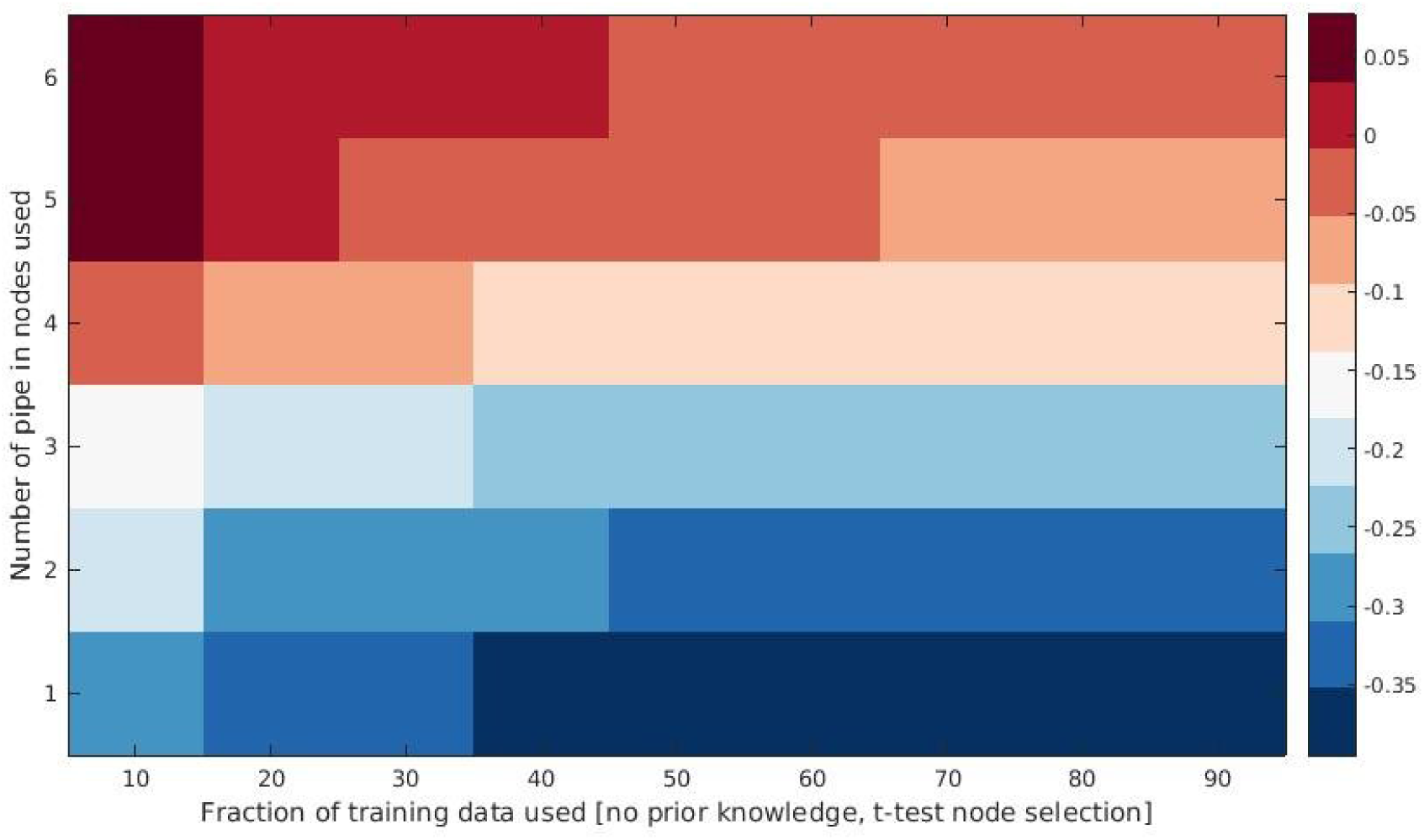
The value of prior knowledge versus the value of additional training data for the baseline networks. The color values indicate (classification accuracy of prior knowledge approach) – (classification accuracy of no-prior knowledge approach). Note that for the prior knowledge runs, 10% training size was used throughout. Full prior knowledge (knowing the six pipe-in nodes in this case) and having a training dataset size of 10% (of the total population size simulated, which is 720 for the base case, thus 72 samples) gives a classification accuracy on par with having five times as much data (50%) and no prior knowledge.

The notion of prior knowledge is a vague one. In our case, the knowledge of which nodes pipe into the nodes we are trying to predict worked well whereas PCA compression of module level steady state data did not. We speculate that pathway-level data compression has been shown to be effective for real genomic datasets [13, 14, 15, 16, 17] because the pathways chosen are intrinsically selecting the genes known to show errant behavior in cancer, for example. In theoretical studies like ours, and in moving this type of thinking to practice, it will be interesting to further investigate various forms of prior knowledge, both more complete (for example, knowledge of state equations) and less complete (less certainty about the pipe-in nodes, or networks where the super-node wiring is more complex). We make the datasets and code to generate them freely available for researchers to test their methods on, which we hypothesize will only strengthen our claim of the value of prior knowledge.

While we considered constructing our networks to mimic known signaling and metabolic pathways in cancer [18], we opted for random networks to ensure hard learning problems and to avoid only representing canonical cancer pathways. Nevertheless, we think it would be interesting to try learning algorithms on curated Boolean and non-Boolean cancer network reconstructions, which there are many of, e.g.http://cellcollective.org. Fumiã *et al* [18] show that different initial state vectors lead to qualitively different phenotypic states (attractors) of their networks (apoptotic – active caspases, immortalized – active hTERT, migratory – inactive E cadherin, etc.). We ran the same networks from different initial conditions and did not observe such initial state dependent behavior, likely due to the random rules and wiring of our networks as opposed to networks tuned for distinct functions and tuned over millions of years of evolution.

Random forest and non-linear support vector machines are known to be able to learn complex rules, including Boolean logic [19]. Since biological systems have inherent Boolean logic embedded in them (consider the need for the use of IF, AND, OR, NOT, etc. in the description of any biochemical process) [20], improved prediction of their behavior will require methods that can represent Boolean logic. RF and SVM methods outperform the more standard statistical learning techniques, including lasso and elastic net, that do not consider complex interactions amongst the input variables, Figure 4. Clustering methods are also often used to analyze genomic data, but our results indicate that Boolean logic is better learned with RF and SVM methods. Of course, if enough data were available such that all sample types were represented, clustering methods could become a workable choice. Neural network methods, including deep learning architectures, thrive in situations where training data is abundant. Given that the main problem that we are working towards, predicting if a particular drug will be effective for a particular patient, does not fall into this situation, we have opted for RFs and SVMs in order to keep the focus on the value of prior knowledge, which we believe – and as evidenced by the results herein – will provide much more sizeable gains for the prediction problem than the particular choice of the non-linear machine learning algorithm used. However, if massive amounts of training data become available, deep learning methods should be assessed as well.

We show that algorithms that are able to find combinatorial patterns are aided by pre-selection of important variable using non-combinatorial techniques. Choosing network nodes which independently have a high predictive value for the learning problem (assessed by univariate t-test, mutual information, or chi-squared methods, which all produced similar results) and using only these nodes in the RF algorithm increases the overall predictive capability, Figure 4. However, univariate selection, since it fails to select all the pipe-in nodes, cannot compete with prior knowledge. We see that t-test selection often selects nodes that, while apparently useful for the classification problem, are likely spurious correlations, whereas the nodes that are actually useful for classification, the pipe-in nodes, must often be only useful in combination with the rest of the pipe-in nodes, and thus not ranked highly by the t-test approach.

While our main result is on the usefulness of prior knowledge, we also study how the difficulty of the learning problem depends on network parameters, notably network size, mutation rate, and number of pipe-in nodes. The trend that we observe is that as these parameters increase, the problem gets harder, but for all three of these there is a point after which the problem seems to get easier. This result, while statistically significant regarding a p-test, is less so if interpreted with the CLES score. Nonetheless, examining the underlying data shows that as mutation rate and number of pipe-in nodes increased past the baseline values, the variance in the super-node values to predict decreases (there is less of a 50-50 split), which may explain the easier learning problem. We observe the same dip when we analyze the data using AUC instead of CA.

Many custom algorithms have been written to interpret and classify genetic data, including approaches using traditional techniques such as logistic regression and attempts using more modern methods, for example matrix factorization as reviewed in [21]. The most relevant to our work are methods which utilize the underlying network structure of the data, such as [22, 23], however these methods make Gaussian assumptions which we wanted to avoid, and these methods are also unsupervised, which is not the problem we study. More importantly, none of the methods to our knowledge use or assess the value of prior knowledge (with the exception of methods that use gene sets and pathways, which is an implicit form of prior knowledge, as already discussed), the main contribution of our work. Assuming that predicting the behavior of complex Boolean networks provides an analogy for predicting biological systems, this paper gives a strong indication that incorporating information about the system being learned in the machine learning process will yield substantial accuracy improvements.

## Acknowledgement

The authors thank Josh Reed for helpful discussions and explorations of the various methods we used.

## References

[1] Matthew Holderfield, Marian M Deuker, Frank McCormick, and Martin McMahon. Targeting RAF kinases for cancer therapy: BRAF mutated melanoma and beyond. Nature reviews. Cancer, 14(7): 455, 2014.

[2] Nicholas McGranahan and Charles Swanton. Biological and therapeutic impact of intratumor heterogeneity in cancer evolution. Cancer cell, 27(1): 15–26, 2015.

[3] Min Huang, Aijun Shen, Jian Ding, and Meiyu Geng. Molecularly targeted cancer therapy: some lessons from the past decade. Trends in pharmacological sciences, 35(1): 41–50, 2014.

[4] Bert Vogelstein, Nickolas Papadopoulos, Victor E Velculescu, Shibin Zhou, Luis A Diaz, and Kenneth W Kinzler. Cancer genome landscapes. science, 339(6127):1546–1558, 2013.

[5] Sweta Mishra and Johnathan R Whetstine. Different facets of copy number changes: permanent, transient, and adaptive. Molecular and cellular biology, 36(7): 1050–1063, 2016.

[6] JG Liao and Khew-Voon Chin. Logistic regression for disease classification using microarray data: model selection in a large p and small n case. Bioinformatics, 23(15): 1945–1951, 2007.

[7] Anastasia Chalkidou, Michael J ODoherty, and Paul K Marsden. False discovery rates in PET and CT studies with texture features: a systematic review. PloS one, 10(5):e0124165, 2015.

[8] L. Raeymaekers. Dynamics of boolean networks controlled by biologically meaningful functions. Journal of Theoretical Biology, 218(3): 331–341, 2002.

[9] I. Shmulevich, E. Dougherty, S. Kim, and W. Zhang. Probabilistic boolean networks: a rule-based uncertainty model for gene regulatory networks. Bioinformatics, 18(2): 261–274, 2002.

[10] Albert-Laszlo Barabasi and Zoltan N Oltvai. Network biology: understanding the cell’s functional organization. Nature reviews genetics, 5(2): 101–113, 2004.

[11] Gail M Sullivan and Richard Feinn. Using effect size-or why the P value is not enough. Journal of graduate medical education, 4(3): 279–282, 2012.

[12] Kenneth O McGraw and SP Wong. A common language effect size statistic. Psychological bulletin, 111(2): 361, 1992.

[13] Michael R Young and David L Craft. Pathway-informed classification system (pics) for cancer analysis using gene expression data. Cancer Informatics, 15:151, 2016.

[14] Y. Drier, M. Sheffer, and E. Domany. Pathway-based personalized analysis of cancer. Proceedings of the National Academy of Sciences, 110(16): 6388–6393, 2013.

[15] C. Vaske, S. Benz, J. Sanborn, D. Earl, C. Szeto, J. Zhu, D. Haussler, and J. Stuart. Inference of patient-specific pathway activities from multi-dimensional cancer genomics data using PARADIGM. Bioinformatics, 26(12): i237–i245, 2010.

[16] A. Tarca, S. Draghici, P. Khatri, S. Hassan, P. Mittal, J. Kim, C. Kim, J. Kusanovic, and R. Romero. A novel signaling pathway impact analysis. Bioinformatics, 25(1): 75–82, 2009.

[17] Shinuk Kim, Mark Kon, and Charles DeLisi. Pathway-based classification of cancer subtypes. Biology Direct, 7(1): 21, 2012.

[18] Herman F Fumiã and Marcelo L Martins. Boolean network model for cancer pathways: predicting carcinogenesis and targeted therapy outcomes. PloS one, 8(7):e69008, 2013.

[19] Ken Sadohara. Learning of boolean functions using support vector machines. In International Conference on Algorithmic Learning Theory, pages 106–118. Springer, 2001.

[20] Rui-Sheng Wang, Assieh Saadatpour, and Reka Albert. Boolean modeling in systems biology: an overview of methodology and applications. Physical biology, 9(5): 055001, 2012.

[21] Andrew V Kossenkov and Michael F Ochs. Matrix factorisation methods applied in microarray data analysis. International journal of data mining and bioinformatics, 4(1): 72–90, 2010.

[22] Safiye Celik, Benjamin A Logsdon, and Su-In Lee. Efficient dimensionality reduction for high-dimensional network estimation. In ICML, pages 1953–1961, 2014.

[23] Safiye Celik, Benjamin A Logsdon, Stephanie Battle, Charles W Drescher, Mara Rendi, R David Hawkins, and Su-In Lee. Extracting a low-dimensional description of multiple gene expression datasets reveals a potential driver for tumor-associated stroma in ovarian cancer. Genome Medicine, 8(1): 1, 2016.

